# matOptimize: A parallel tree optimization method enables online phylogenetics for SARS-CoV-2

**DOI:** 10.1101/2022.01.12.475688

**Authors:** Cheng Ye, Bryan Thornlow, Angie Hinrichs, Devika Torvi, Robert Lanfear, Russell Corbett-Detig, Yatish Turakhia

## Abstract

Phylogenetic tree optimization is necessary for precise analysis of evolutionary and transmission dynamics, but existing tools are inadequate for handling the scale and pace of data produced during the COVID-19 pandemic. One transformative approach, online phylogenetics, aims to incrementally add samples to an ever-growing phylogeny, but there are no previously-existing approaches that can efficiently optimize this vast phylogeny under the time constraints of the pandemic. Here, we present matOptimize, a fast and memory-efficient phylogenetic tree optimization tool based on parsimony that can be parallelized across multiple CPU threads and nodes, and provides orders of magnitude improvement in runtime and peak memory usage compared to existing state-of-the-art methods. We have developed this method particularly to address the pressing need during the COVID-19 pandemic for daily maintenance and optimization of a comprehensive SARS-CoV-2 phylogeny. Thus, our approach addresses an important need for daily maintenance and refinement of a comprehensive SARS-CoV-2 phylogeny.

**Significance Statement:** Phylogenetic trees have been central to genomic surveillance, epidemiology, and contact tracing efforts during the COVD-19 pandemic. With over 6 million SARS-CoV-2 genome sequences now available, maintaining an accurate, comprehensive phylogenetic tree of all available SARS-CoV-2 sequences is becoming computationally infeasible with existing software, but is essential for getting a detailed picture of the virus’ evolution and transmission. Our novel phylogenetic software, matOptimize, is helping refine possibly the largest-ever phylogenetic tree, containing millions of SARS-CoV-2 sequences, thus providing an unprecedented resolution for studying the pathogen’s evolutionary and transmission dynamics.

## Introduction

With over 6 million genome sequences now available on online databases, SARS-CoV-2 is the most sequenced pathogen in history by far (1). Phylogenetics has been a foundational tool in analyzing this vast volume of genomic data for a public health response during the COVID-19 pandemic (2). For example, phylogenetics has been instrumental in genomic surveillance, i.e. for identifying and naming its new variants (Rambaut et al., 2020) and for tracking the different SARS-CoV-2 variants circulating in a given geographic region (3, 4). Phylogenetic trees have also helped in establishing transmission links between infections (5), in disambiguating community transmission from outside introductions (6, 7), in identifying the mutations that might have conferred increased transmissibility to the virus (8, 9), and for estimating the reproduction number (R0) of the virus and its variants (10, 11).

Many of these far-reaching applications benefit from having a comprehensive phylogenetic tree – one produced without sub-sampling the available sequences. For example, when all available SARS-CoV-2 sequences are not represented in a single phylogenetic tree, transmission links between isolates may be lost. Likewise, sub-sampled phylogenetic trees could omit important lineages or sub-lineages corresponding to different variants of the virus. This can have adverse consequences on the downstream evolutionary and epidemiological studies. A comprehensive and up-to-date phylogenetic tree is therefore an essential goal for the global health response to the SARS-CoV-2 pandemic.

Online phylogenetics is a potentially transformative solution for the massive computational challenges imposed by continual genome sequence collection during the pandemic (12, 13). It involves incorporating new sequences as they become available onto an existing phylogeny and periodically refining the updated phylogeny. The recent development of UShER (14), which uses stepwise addition of new sequences onto an existing phylogenetic tree, helped overcome an important computational barrier for the maintenance of a comprehensive SARS-CoV-2 phylogenetic tree (15). For large SARS-CoV-2 phylogenies, UShER’s maximum parsimony (MP) approach works exceedingly well, and may even be tantamount to the far more computationally-expensive maximum likelihood (ML) approaches available currently (16). However, since UShER is based on the greedy approach of stepwise addition of new sequences, it sometimes places new sequences suboptimally in the tree (14, 17). Tree optimization tools, which use tree rearrangement to find more optimal tree configurations, can help ameliorate this issue. However, in the recent six months (May-October 2021), the UShER-derived comprehensive SARS-CoV-2 phylogenetic tree has more than tripled in size, viz. an average rate of approximately 1.2% additional sequences being incorporated each day. Approaches capable of daily tree optimization of the massive SARS-CoV-2 phylogeny are therefore an urgent necessity for emerging online phylogenetic toolkits.

Phylogenetic tree optimization has been extensively studied in the context of maximum parsimony, and tools such as TNT (18), PAUP* (19), MPBoot (20), PHYLIP (21) and MEGA (22) are already available. These tools take as input an existing phylogenetic tree in the Newick format and the multiple sequence alignment (MSA) of the phylogenetic tips in FASTA format, then compute the parsimonious state assignments for every node of the tree for each alignment site and then use tree rearrangement to find a more parsimonious tree. They also maintain multiple equally parsimonious candidate trees and sometimes use tree drifting (18) during the optimization to avoid getting stuck in local optima. TNT also provides additional heuristics, such as sectorial search (23), to speed up the search. Though these tools are remarkably efficient for the typical applications that they were developed for (such as for one-time species tree inference), they are inadequate for handling the scale and pace of the rapidly expanding SARS-CoV-2 data due to their prohibitive runtime and memory requirements.

Here, we developed matOptimize, a novel MP-based tree optimization tool that uses several domain-specific optimization techniques to greatly speed up, and reduce the cost of, optimizing the comprehensive SARS-CoV-2 phylogeny. Starting from an UShER-derived SARS-CoV-2 phylogeny, matOptimize provides roughly the same parsimony score improvement (used as a measure of tree quality) as the previous state-of-the-art for a moderately-sized phylogeny, and is the only tool that meets the constraints of daily-optimization for the current scale of comprehensive SARS-CoV-2 phylogeny. matOptimize uses memory-efficient data structures to handle the vast scale of SARS-CoV-2 data, and is efficiently parallelized for multicore CPU as well as a high-performance computing (HPC) cluster. This means that the resources allocated to matOptimize can be scaled to meet the computational requirements of the ever-expanding comprehensive SARS-CoV-2 phylogeny. Our analysis on real data suggests that matOptimize indeed qualitatively improves the comprehensive SARS-CoV-2 phylogeny.

## Results

### matOptimize: Algorithm Overview

matOptimize is a parallel algorithm (Figure 1, Methods) for parsimony-based optimization of large SARS-CoV-2 phylogenies. matOptimize takes as input a starting phylogeny, such as the UShER-derived mutation-annotated tree (MAT) (which is a file format that stores a phylogenetic tree in which the branches are annotated with the mutations that are inferred to have occurred on them (14)) or a Newick file, and an optional VCF file specifying the genotypes at the tips of the phylogeny. When a VCF input is provided, matOptimize uses a parallel implementation of the Fitch-Sankoff algorithm (24, 25) to infer the most parsimonious state for every VCF site at every branch of the tree. Otherwise, the algorithm accepts the states annotated in the MAT and skips to the next step. matOptimize uses a modified MAT (Figure 1B,C) for a highly memory-efficient storage of the states (such as the Fitch states (24), see Methods) internal to the tree that are needed for optimizing the tree. Next, matOptimize begins the first iteration of the parallel search phase for exploring tree rearrangement, using subtree pruning and grafting (SPR) moves, starting from all possible source nodes in the tree. matOptimize uses the incremental update method of Gladstein et al. (26) (Methods, Figure 1B,C), modified slightly to handle polytomies, to quickly discover profitable tree moves. The search for profitable moves is parallelized across multiple available CPU nodes using MPI (https://www.mpich.org/). The tree object is held immutable throughout this parallel search phase in matOptimize, so that each process can explore different tree rearrangements independently, with minimal inter-process communication.

**Figure 1.**
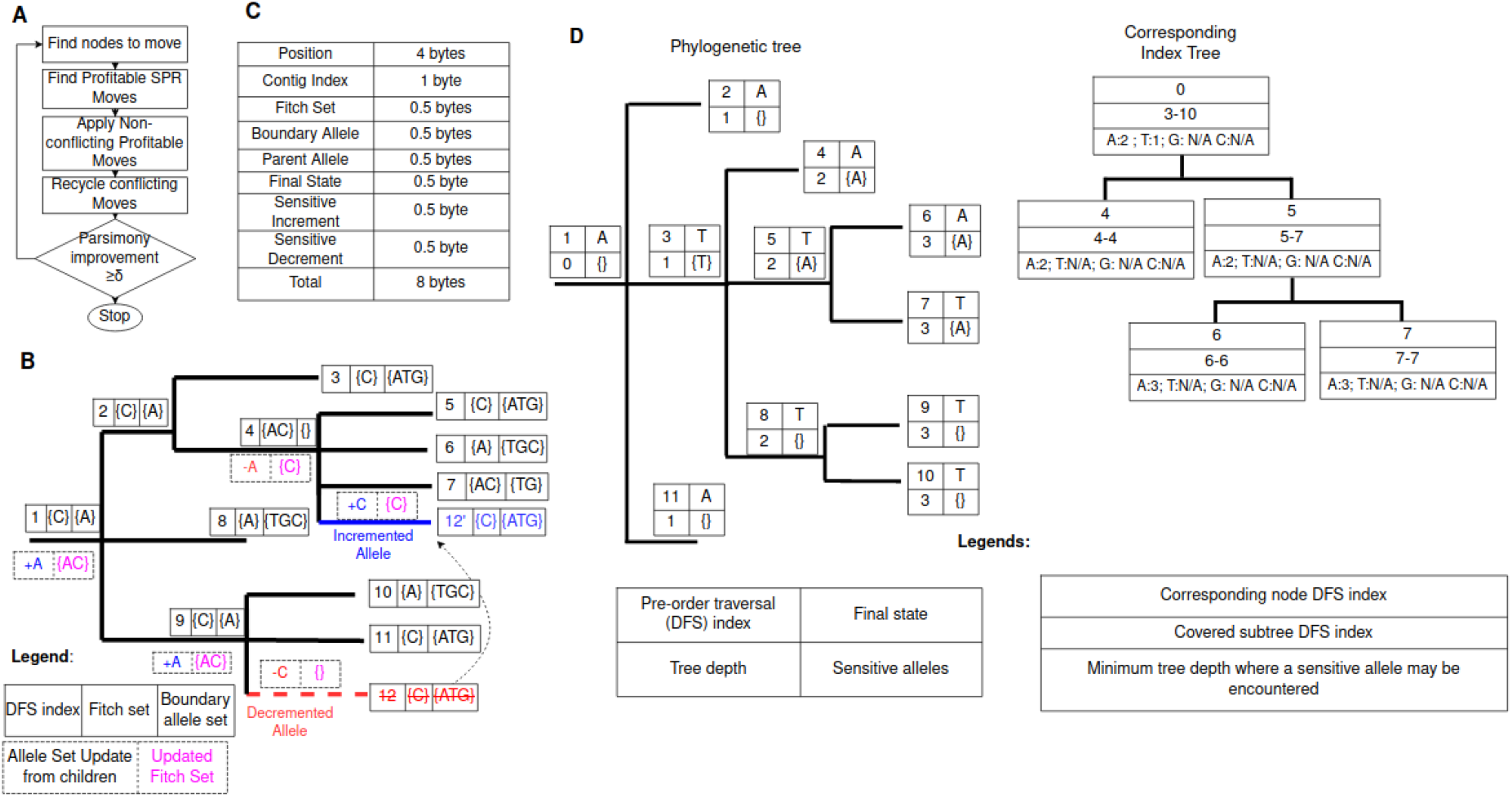
An illustration of the matOptimize algorithm. (**A)** A flowchart of the different algorithmic stages in matOptimize. (**B**) An example of how matOptimize estimates the parsimony score improvement achievable from a single SPR move using a small number of steps without redoing the entire Fitch algorithm (24). Each node of this tree has an integer label (1 to 12) and is annotated with its Fitch and boundary allele sets (Methods). This example evaluates a move in which the subtree rooted at node 12 is pruned from node 9 and regrafted at node 4, and the corresponding updates to the Fitch sets of the affected nodes and their ancestors. (**C)** Storage requirements for a mutation in the mutation-annotated tree (MAT) data structure of matOptimize. This is a modified version of the original MAT proposed in UShER (14) in order to maintain auxiliary information (such as Fitch and boundary allele sets) for performing optimization. Each mutation in matOptimize is stored compactly using only 8 bytes, which helps it maintain a small memory footprint overall. (**D**) An example phylogenetic tree (left) and its corresponding index tree (right). The index tree is used to speed up the search for promising destination nodes for SPR moves from a single source node via search space pruning (Methods). Each node in the phylogenetic tree is annotated with (i) a pre-order traversal index, (ii) the depth of the node, (iii) the final allele assignment and (iv) the sensitive allele (Methods). The index tree uses a B-Tree (27) to store the nodes at which the sensitive alleles are found (Methods). Each node in the index tree corresponds to one node in the phylogenetic tree and stores the DFS index range of the subtree that the node covers and the minimum depth within the subtree at which a sensitive allele may be encountered. “N/A” implies that the allele is not present in the subtree.

An iteration begins with the main process distributing source nodes to worker processes. Then, the worker processes evaluate (without applying) all possible SPR moves from the assigned source nodes and periodically return the profitable moves to the main process, which applies non-conflicting moves to update the tree and begin the next iteration. During the parallel search phase, each process can take advantage of the parallelism available on each compute node by distributing the tree rearrangements to be explored to multiple threads using Intel’s TBB library (https://github.com/oneapi-src/oneTBB, Methods). Since most parsimony score improvements come from small SPR radius moves which are quick to evaluate (Figure S1), matOptimize starts with a small maximum SPR radius of 2, which is doubled at the end of each iteration till the maximum possible tree radius (twice the tree depth) is reached.

### matOptimize rapidly and efficiently optimizes massive SARS-CoV-2 phylogenies

We used the database of UShER-based SARS-CoV-2 trees (15) to pick phylogenies at three different time points, containing 100K, 1M and 3M samples, respectively. We used these as starting phylogenies to compare the performance of matOptimize with TNT (28), the fastest and most efficient MP-based tree optimization tool currently available (Figure 2, Methods). Since TNT is not multithreaded, it had to be manually parallelized across multiple processes (Methods). We also limited the optimization to 24 hours to meet the constraints for enabling daily tree optimization which is required for comprehensive phylogenetics during the pandemic. Of the three trees we evaluated, TNT (23) completed its optimization for only one, i.e., the smallest 100K-sample tree, achieving a parsimony score improvement of 0.63% in 1.13 hours. matOptimize completed this tree optimization 19x faster and with 37x lower peak memory (Figure 2A), achieving a comparable but slightly smaller parsimony improvement of 0.59%. The smaller parsimony score improvement of matOptimize could be explained by the fact that it searches from one starting tree and uses SPR moves only, which have a smaller search space than the TBR moves and the independent search of multiple trees used in TNT (29).

**Figure 2:**
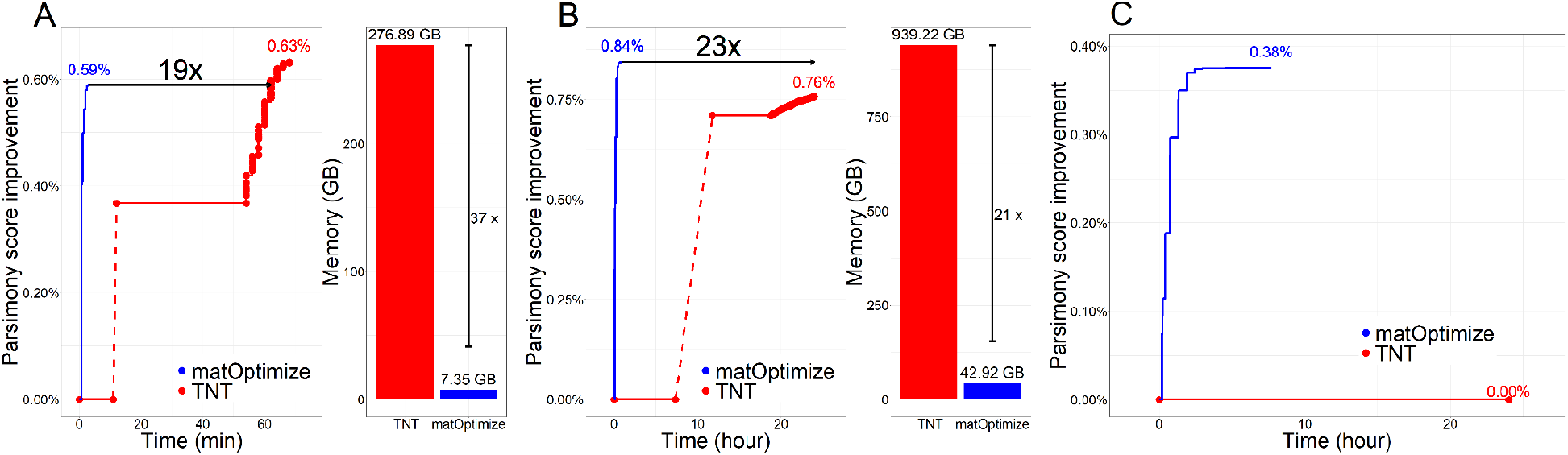
Comparison of parsimony score improvement and peak memory requirement of matOptimize and TNT starting from an SARS-CoV-2 based UShER-derived (**A**) 100K-sample tree, (**B**) 1M-sample tree, and (**C**) 3M-sample tree. For (C), the peak memory requirement is not shown since TNT did not begin the optimization phase by the time it was terminated after 24 hours of execution.

The benefits of matOptimize become apparent when the tree size gets larger – it completed optimization for the 3M-sample tree in only 7.7 hours (Figure 2C). On this tree, TNT did not even start applying profitable moves. This is because, for the scale of this tree, TNT spent the entire day loading the input files and initializing its internal data structures. Since matOptimize also uses SPR radius doubling at each iteration, and since smaller radii iterations are quick to complete, the largest improvements in parsimony score are achieved very quickly during optimization (Figure 2, S1). Due to its memory-efficient internal data structures and parallel implementation of the Fitch-Sankoff algorithm during initialization (Methods and Discussion), matOptimize starts optimizing the tree after a small delay, which is desirable for time-constrained optimization scenarios imposed by the pandemic. In contrast, TNT incurs a long startup delay followed by a slow improvement of parsimony. matOptimize is therefore especially well suited for SARS-CoV-2 online phylogenetics (13).

Detailed analysis of the resulting tree revealed that matOptimize indeed improves the tree quality. For example, on the 100K-sample tree, the 0.59% parsimony score improvement also translated into a 0.48% improvement in the log likelihood score – from −919354.028 to −914925.186 (Methods). This is consistent with our previous analyses (13) where we observed that parsimony and likelihood scores are strongly correlated for SARS-CoV-2 phylogenies. We also observed that in some cases, the PangoLEARN lineage assignments (30) appeared a little less dispersed through the tree (Figure S2), with the Pango lineage parsimony score decreasing from 17043 to 17014 post optimization (Methods). Here, the small improvement through optimization reaffirms the observation that even the stepwise addition of samples through UShER has high accuracy for lineage assignments (15, 31).

### matOptimize scales well in runtime and memory requirement with the number of available processors

Efficient parallelism ensures that matOptimize can keep up with the increasing computational demands for optimizing SARS-CoV-2 phylogenies. We designed it to parallelize over all available virtual cores on a CPU (vCPUs), as well as a large number of CPUs available in a high-performance computing (HPC) cluster, which includes cloud platforms. Figure 3A shows that matOptimize shows a high strong scaling efficiency. matOptimize requires 109 minutes to complete one round of optimization on the 1M-sample tree with 64 vCPUs over two CPU instances, which reduces to 11.5 minutes with 1024 vCPUs over 32 CPU instances (scaling efficiency of 59%, Figure 3A). In the latter, the parallel search for profitable moves required 7.5 minutes and the sequential step of applying the moves took 4 minutes. We therefore expect strong scaling efficiency to improve as the SARS-CoV-2 tree gets bigger, and the parallel search phase consumes a larger fraction of the total runtime.

**Figure 3:**
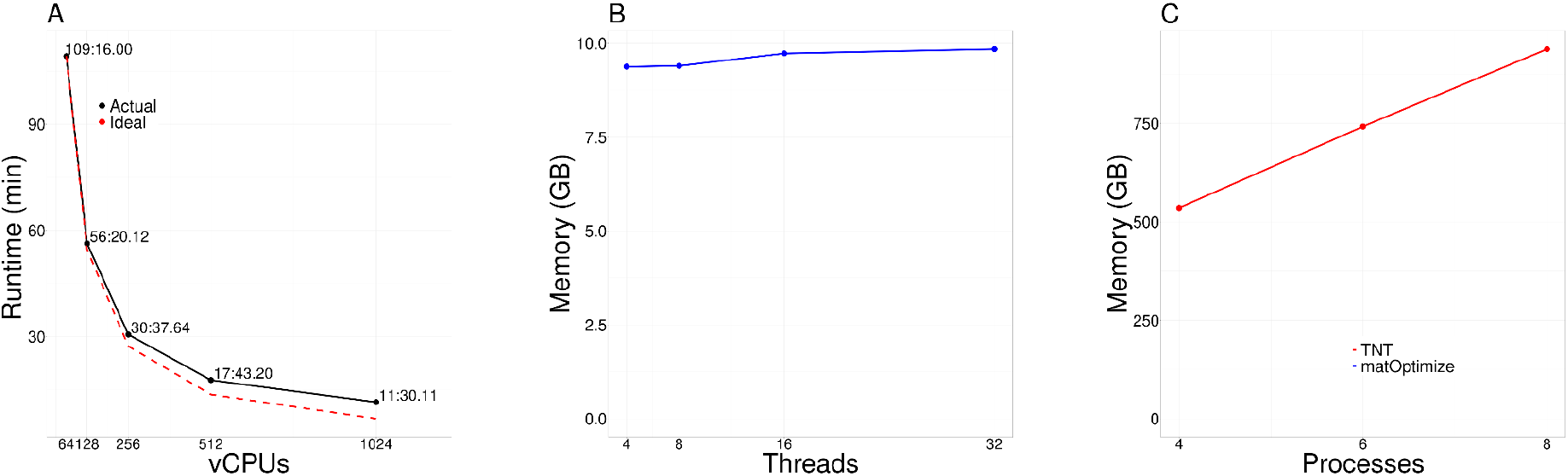
Performance and memory scaling efficiency of matOptimize using the 1M-sample tree. (**A**) Strong multi-node scaling efficiency of matOptimize. Each node is a Google Cloud Platform (GCP) instance e2-highcpu-32 consisting of 32 vCPUs and 32 GB memory. The number above each data point corresponds to the actual runtime in minutes:seconds format. (**B**) The peak memory requirement of matOptimize is small (below 10 GB) and remains roughly constant with the number of CPU threads. This allows matOptimize to exploit all available parallelism on a multicore CPU instance without being limited by the available memory. (**C**) In comparison, the peak memory requirement of TNT is large (>500 GB) and increases linearly as the parallelism is increased. This limits the amount of parallelism that TNT can exploit – in the example shown, TNT could exploit only up to 8 available vCPUs out of the 40 available on the memory-optimized GCP instance m1-ultramem-40 before running out of memory.

A key reason why matOptimize scales efficiently is its low memory requirement. This is enabled through its compact mutation-annotated tree data structure derived from UShER, which we use in lieu of the traditional matrix representation of the MSA in existing frameworks (Figure 3B, 1C, Methods). When the number of CPU threads used by matOptimize is increased from 4 to 32, the memory requirement for the 1M-sample SARS-CoV-2 tree increases only marginally, from 9.38 GB to 9.84 GB (Figure 3B). In contrast, the large and linear memory requirement of TNT limits the amount of parallelism it can exploit (Figure 3C). For example, with the 1M-sample SARS-CoV-2 tree, we could only parallelize TNT to utilize 8 of the 40 available vCPUs on the m1-ultramem-40 instance that we used on the Google Cloud Platform (GCP), since eight TNT processes consumed nearly all of 961 GB available memory on this instance. It is therefore unlikely that TNT would scale to the much larger SARS-CoV-2 phylogenies anticipated in the future due to its prohibitive memory requirement.

### Larger SPR radii provide smaller, but measurable improvements in parsimony score

We explored the contribution of different SPR radii to the improvement in parsimony score on the 100K-sample tree, 1M-sample tree, and the 3M-sample tree (Figure S1). On the 100K-sample tree, a small SPR radius of 2 alone could achieve 70% of the total parsimony score improvement. For the 3M-sample tree, SPR moves of radius 2 could achieve only 20% of the total possible parsimony score improvement. However, even for larger trees, the parsimony score still improves most rapidly with small radii (Figure S1) – 90% of parsimony improvement is achieved with less than 25% of the total runtime for both the 1M-sample and 3M-sample trees. Therefore, matOptimize doubles the search radius after each round in order to provide the largest parsimony improvement at the early stages of optimization. The difference between radius doubling and fixed large radius strategy is negligible (Supplementary Table 1). Combined with the small preprocessing time, this radius doubling strategy makes matOptimize particularly suitable for “online” optimization of large SARS-CoV-2 phylogenies. The reason is that it may be necessary to halt optimization if the runtime exceeds the time allowed until additional sample additions are necessary.

### Multi-node parallelization of Phylogenetic Placement

Since tens of thousands of new SARS-CoV-2 genome sequences are being shared through online databases every day over and above the millions that already exist (32), even the daily stepwise incorporation of new sequences using UShER (14) has also become challenging. Therefore, we also accelerated the new bottleneck step of phylogenetic placement through multi-node parallelization. To do this, we developed a new utility, called *matUtils merge*, that can merge two input mutation-annotated trees (MATs) that have a consistent backbone tree. matUtils merge operates by first identifying the correspondence between the shared nodes of the two input MATs and then placing unique samples in one MAT onto the other MAT by restricting the placement search to a small radius around the nearest common node, which results in a significant speedup over an exhaustive tree placement search (14). With matUtils merge, we split up the VCF containing new samples uniformly and distribute them over independent CPU nodes, each executing UShER to place the corresponding samples on the same base mutation-annotated tree (MAT). The resulting MAT files are then merged into a single output MAT containing all new samples using a parallel reduction tree of a new utility called *matUtils merge* (Figure S3). With this multi-node parallelization, UShER could place 100K new samples on the 1M-sample tree in under 38 minutes with 1024 vCPUs, achieving 11.6x speedup over 64 vCPUs (Figure S4). While we found the distributed placement approach used in matUtils merge to be slightly suboptimal (i.e., resulting in a higher parsimony score) compared to a completely sequential approach of UShER, we found no noticeable difference in the resulting trees following matOptimize optimization (Figure S5). A combined pipeline using UShER, matUtils merge and matOptimize is currently being used by UCSC Genome Browser to maintain on a daily basis and share a comprehensive SARS-CoV-2 phylogeny (https://genome.ucsc.edu/cgi-bin/hgPhyloPlace).

## Discussion

Several evolutionary studies, such as inferring SARS-CoV-2 recombination (33) or characterizing the transmissibility of different mutations (9), rely on, or can benefit from, the knowledge of a comprehensive SARS-CoV-2 phylogenetic tree. Currently, the UCSC Genome Browser uses a comprehensive SARS-CoV-2 phylogenetic tree which is updated daily through phylogenetic placements of new sequences performed by UShER (34). This tree is being used widely by health officials and researchers worldwide through UShER’s command line (https://github.com/yatisht/usher) and web interface (https://genome.ucsc.edu/cgi-bin/hgPhyloPlace) to perform genomic surveillance (31, 35–37) and contact tracing (5) of the COVID-19 pathogen. This tree has also been applied for SARS-CoV-2 lineage assignment in the optional mode of Pangolin (https://github.com/cov-lineages/pangolin/), the most widely used scientific nomenclature system for SARS-CoV-2 lineages (38), as well as in several recently developed phylogenetic toolkits (7, 39, 40). GISAID’s Audacity tree (1) is also based on the UShER package. Maintaining a high-quality comprehensive SARS-CoV-2 phylogeny consisting of all available SARS-CoV-2 sequences has emerged as a problem of utmost importance during the COVID-19 pandemic.

Currently, matOptimize appears to be the only viable tree optimization tool for SARS-CoV-2 online phylogenetics. matOptimize’s performance is the result of several optimization and parallelization techniques that it incorporates. For example, matOptimize benefits from the recently developed, space-efficient data structure of a mutation-annotated tree (MAT) – an uncompressed MAT file of 3 million SARS-CoV-2 sequence requires only 136 MB to encode basically the same information that is contained in an 88 GB multiple-sequence alignment (MSA) FASTA file. By using a modified MAT, matOptimize can compactly store all the information necessary for optimization and avoid the high cost of maintaining the entire alignment in memory. Widely-used phylogenetic programs, such as TNT (28), MPBoot (20) and MEGA (22), use strategies for parsimony score optimization that are far too expensive at SARS-CoV-2-scale. As a result of its scale up (more vCPUs per node) and scale out (more CPU nodes) capability, matOptimize is the only parsimony-based tree optimization software that is currently able to meet the computational demands for optimizing the ever-increasing SARS-CoV-2 phylogeny. Overall, matOptimize is orders of magnitude more cost and memory efficient than the state-of-the-art, and we expect that matOptimize would help maintain a high-quality comprehensive SARS-CoV-2 phylogeny for several months to come and for several other densely sampled pathogens as well in the future.

## Methods

### Material

We collected SARS-CoV-2 genome sequence data from major online databases: GISAID (1), COG-UK (41), GenBank (42) and CNCB (https://bigd.big.ac.cn/ncov/release_genome) until August 25, 2021 and used it to build a comprehensive SARS-CoV-2 phylogeny based on the methodology described in (15). We used the GenBank MN908947.3 (RefSeq NC_045512.2) sequence as the reference for rooting the tree and used the sampling date metadata to derive from our comprehensive tree three subtrees containing the earliest 100K, 1M and 3M samples, referred to as 100K-sample tree, 1M-sample tree, and 3M-sample tree, respectively.

### Algorithm description

Given a starting MAT file or a combination of a starting tree in Newick format along with the genotypes of its tips (samples) in VCF format, matOptimize aims to reduce the parsimony score of the tree using subtree pruning and regrafting (SPR) moves. matOptimize allows the VCF file to specify ambiguous bases using the International Union of Pure and Applied Chemistry (IUPAC) format (https://www.bioinformatics.org/sms/iupac.html), and the ambiguity is maintained throughout the optimization using a modified mutation-annotated tree format (Figure 1C).

Figure 1 describes the matOptimize algorithm and data structures. The matOptimize algorithm performs several iterations of optimization consisting of two phases: (1) the parallel search phase and (2) the non-conflicting moves application phase, described in more detail below. The iterations continue until the percent parsimony score improvement in the last iteration is less than the user specified threshold (δin Figure 1A, default: 0.5%), or the wall-clock time limit specified by the user (default: unlimited) is exceeded (Figure 1A). Only one tree is maintained throughout the optimization in matOptimize. We use the following terminologies for the remaining description of the algorithm:

- *Child*: A direct descendant of a node
- *Allele frequency*: The number of children of the node that contains the allele in its Fitch set. On leaf nodes, zero for alleles not observed, 1 for observed alleles.
- *Tree depth*: the distance from root to a node.
- *Fitch set*: The most common allele(s) among the Fitch sets of the children of the current node. On leaf nodes, Fitch sets are observed alleles.
- *Major allele frequency*: Allele frequency of an allele in the Fitch set (the same for all alleles in the Fitch set).
- *Boundary allele set*: Alleles whose frequency is exactly one less than the major allele frequency.
- *Final state*: A single allele inside the major allele set obtained by arbitrary tie-breaking strategy among multiple equally-parsimonious alleles.
- *An allele is incremented*: An allele is added to the Fitch set of a child of a node.
- *An allele is decremented*: An allele is removed from the Fitch set of a child of a node.
- *Least common ancestor (LCA) nodes*: The node furthest from the root that is the ancestor of both source node and destination node.
- *Sensitive Allele*: An allele that will improve the parsimony score of the tree when incremented alone, excluding parent alleles. Parent alleles are excluded because they do not contribute to parsimony scores.
- *DFS*: post order depth first search.

#### 1. Parallel Search Phase

In this phase, all nodes are searched in parallel for SPR moves within a fixed radius that may reduce the parsimony score if applied to the starting tree alone. The SPR radius starts with 2 and doubles at every iteration. To calculate the parsimony score change of a single move, matOptimize uses a variant of Gladstein’s incremental update method (26) that can also handle polytomies. When sample genotypes are specified through an input VCF file, matOptimize first pre-processes the tree to calculate the Fitch set, boundary allele set, and final state for every site at every node of the tree. matOptimize also parallelizes Fitch-Sankoff computations for different loci and parsing of chunks of VCF using Intel’s TBB library (https://github.com/oneapi-src/oneTBB). matOptimize calculates the change in parsimony score from a single move by tracking the change in major allele frequency as the effect of the move propagates toward the root.

An allele can be added to the Fitch set from the boundary allele set (i) if it is incremented, or (ii) if all alleles in the Fitch set are decremented. An allele is removed from the Fitch set when it has been decremented, or if some other alleles in the Fitch set have been incremented. Finally, the parsimony score is calculated as the sum of change in children count minus change in major allele count at each node on the path from source or destination node to root. The starting tree remains immutable and shared among all threads throughout the parallel search phase, so the memory consumption only increases negligibly with the number of threads.

Figure 1B illustrates how the Fitch set update works with an example SPR move in which the subtree rooted at node 12 is pruned from node 9 and regrafted at node 4. For this move, the alleles in the Fitch set of node 12 must be decremented at node 9 (during pruning) and incremented at node 4 (during regrafting). The Fitch set of node 12 consists of a single allele C. Since C is the only allele in the Fitch set of node 9, the alleles from the boundary allele set of node 9 get added to its Fitch set, resulting in an updated Fitch set, {A,C}, with a lower major allele count. The change in Fitch set of node 9 is propagated upwards to its parent, i.e., to root node 1. During the pruning step, the decrement of the major allele count at node 9 has no effect on the parsimony score since it is offset by the decrement in the children count. During regrafting, since allele C, which is already present in the Fitch set of node 4, is incremented, its major allele count is now higher than the remaining alleles in the Fitch set, i.e. allele A, which is decremented. This change is propagated upwards, i.e., to parent node 2, but has no effect on it since allele A is not present in its Fitch set. Since the regrafting step also does not change the parsimony, the net parsimony score change of this move is 0.

#### 2. Search space pruning

To minimize the number of non-profitable moves explored from a source node, matOptimize quickly calculates a lower bound of parsimony score change of moving the pruned subtree to other nodes within a specified radius. It does this by first precomputing the sensitive alleles for each site at each node using a dynamic programming algorithm. The sensitive allele sets are then organized into a B-Tree (43), called index tree (Figure 1D), using the DFS index of the nodes. The internal nodes of the index tree are also annotated with the minimum depth at which each allele will be encountered and corresponding the DFS indices of phylogenetic tree nodes it covers (Figure 1D). During the parallel search phase, the parsimony score change lower bound is initialized with the parsimony score change that would result from removing the source node alone. Then, matOptimize queries the index tree for the minimum depth at which the allele that source node carries may be encountered. If it is larger than the maximum radius minus the distance between the source node and the node being examined, the lower bound of parsimony score change is incremented. In this step, the DFS index range in the index tree is queried to find the minimum depth of the node in the subtree of the currently examined node at which the sensitive allele may be encountered. All destination nodes in a subtree rooted at the node being examined are skipped if the parsimony score change lower bound at this node is higher than that of the most profitable move from the source node found so far (Figure 1D).

#### 3. Multi-node Parallelization Strategy

At the start of the parallel search phase, the main process broadcasts the intermediate MAT to all worker processes, and collects the profitable moves found by worker processes. Worker processes request one minute of workload from the main process, and then the main process replies with one minute of workload or workload for half of the estimated remaining time, whichever is less. Each process may use multiple threads sharing the same tree.

#### 4. Non-conflicting Moves Application phase

Moves that share nodes on the paths from source nodes to destination nodes are defined to be conflicting as they cannot be simultaneously applied. If conflicting moves are found, matOptimize accepts one and defers the remaining to the next iteration. All non-conflicting moves are applied in a batch. matOptimize corrects the Fitch and boundary allele sets of the altered nodes and their ancestors serially in post-order traversal order. Sensitive alleles are recomputed from scratch at the beginning of every new iteration. Profitable source nodes in the last iteration are searched again, until there is no parsimony improvement, after which a new round of iterations is started with doubled SPR radius and all possible source nodes.

### Tree Evaluation

To calculate the likelihood scores of our trees, we used IQ-TREE2 (44) COVID-19 release 2.1.3, with Jukes-Cantor (JC) model and a minimum branch length of 1e-11. Trees were visualized with Taxonium (https://cov2tree.org/). Pango lineages for sequences were assigned using the Pangolin software (30) and parsimony scores for the lineage assignments were computed using TNT (28), taking clade assignment as binary characters.

### Baseline comparison

#### Performance Benchmarking

We used TNT (28) (June 2021 version) as a baseline for comparing performance and peak memory requirements with matOptimize. Given its high peak memory requirements, we used an m1-ultramem-40 instance (40 vCPUs, 961 GB, $6.30/hour) for TNT. For matOptimize, we used 7 e2-highcpu-32 instances for matOptimize that cumulatively have an hourly cost ($5.54/hour) comparable to the m1-ultramem-40 instance. The peak memory requirements were estimated by reading the resident set size (RSS) of each process every two minutes. The parsimony scores were computed after rerooting the trees to GenBank MN908947.3 (RefSeq NC_045512.2).

#### Parallelization of TNT

TNT does not natively include a multithreaded optimization mode, so in order to parallelize it, we used multiple processes performing exclusive sectorial search (XSS), which had better performance compared to parsimony ratchet, tree drifting or direct branch swapping modes in TNT. In the XSS mode, TNT splits the tree into non-overlapping sectors of equal size, and optimizes using each sector separately using global optimization using tree bisection and reconnection (TBR) moves. If the parsimony score improves within the sector, TNT uses it to replace the corresponding sector in the starting tree. We parallelized XSS by splitting the starting tree into 10 non-overlapping sectors, which offered the best runtime and parsimony score tradeoff. During global TBR moves, we let each TNT process optimize the entire tree independently and broadcast the tree to other processes. We parallelized TNT with the highest number of processes possible within the available memory of the instances. This corresponded to 40, 8 and 1 processes for 100K-sample, 1M-sample and 3M-sample trees, respectively.

#### Software Availability

The matOptimize code is available as part of the UShER package (https://github.com/yatisht/usher), which can also be installed via bioconda (https://bioconda.github.io/recipes/usher/README.html). The scripts we used to perform the experiments in this paper are available at https://github.com/yceh/matOptimize-experiments.

## Acknowledgments

We thank Willi Hennig Society for the TNT software. We thank Bui Quang Minh and Diep Thi Hoang for helpful discussions about MPBoot.

C.Y. and Y.T. were supported by the Centers for Disease Control and Prevention grant BAA 200-2021-11554. This work was also supported by the National Institutes of Health grant R35GM128932 (to R.C.-D.). B.T. was supported by National Institutes of Health grants T32HG008345 and F31HG010584. R.L. is supported by an Australian National University Futures Grant, an Australian Research Council Discovery Grant [DP200103151], and a Chan-Zuckerberg Initiative Grant for Essential Open Source Software for Science. This work was also supported by the generous donations by the Eric and Wendy Schmidt Foundation.

## Supplementary Figures And Tables

**Figure S1:**
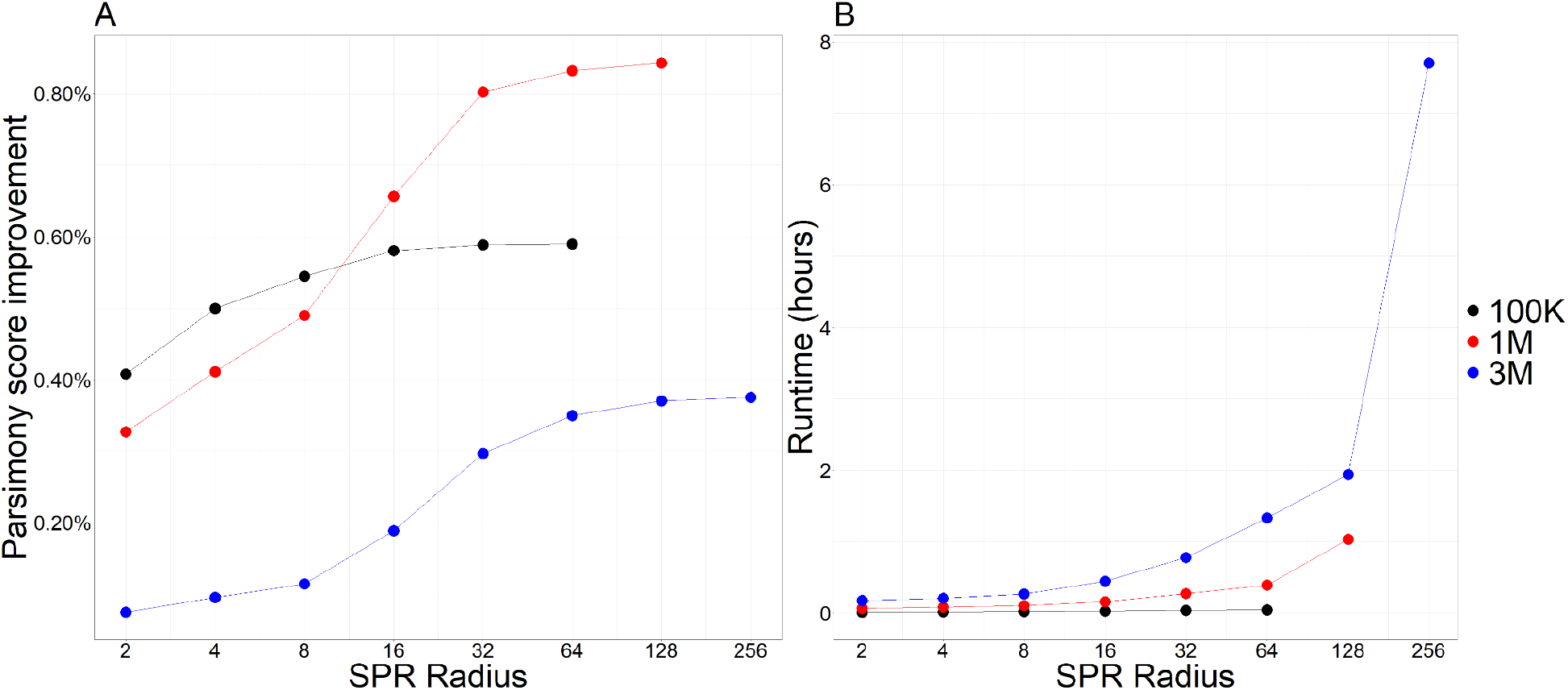
(**A**) Parsimony score improvement and (**B**) the total runtime for different SPR radius achieved through the radius doubling mode in matOptimize.

**Figure S2:**
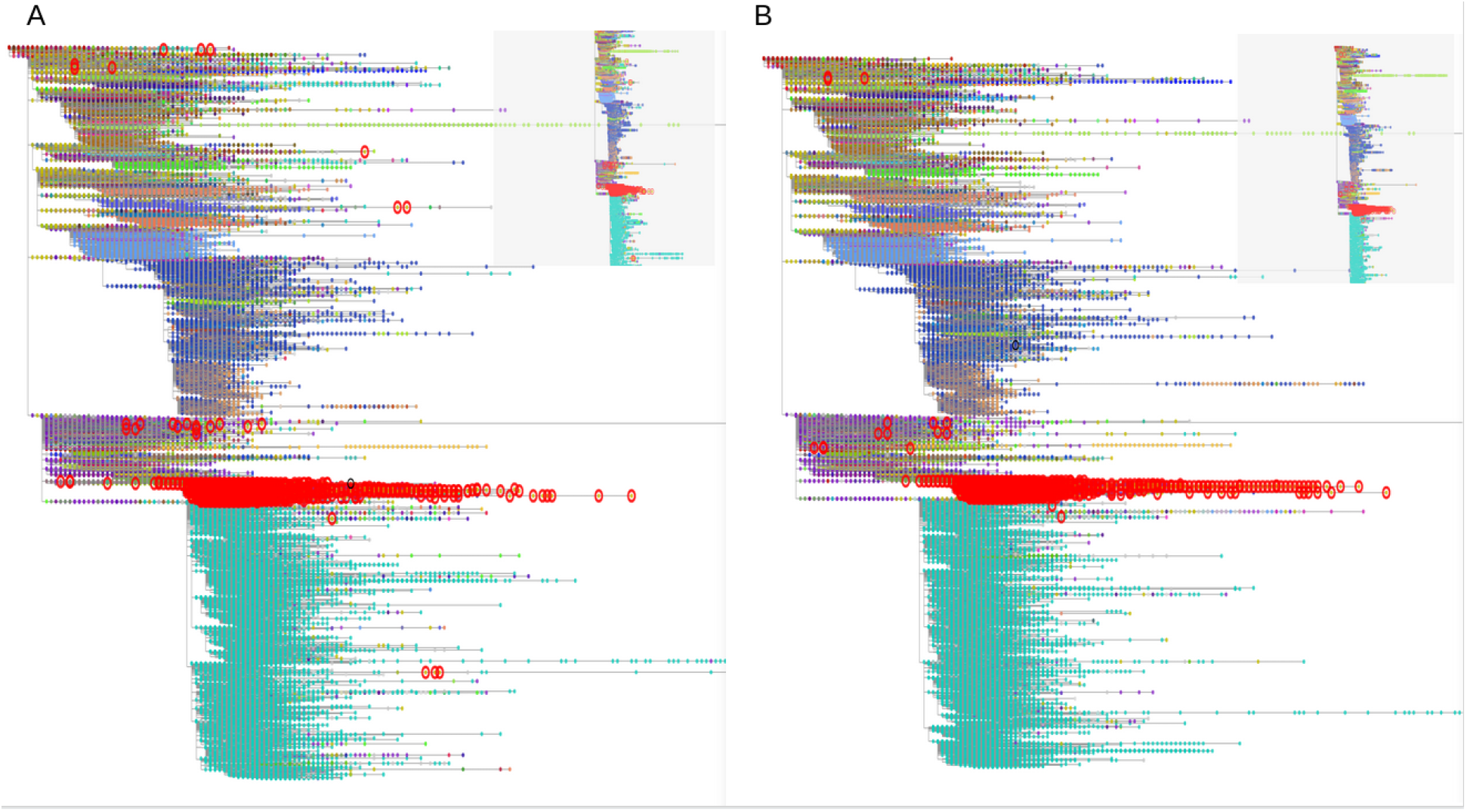
Taxonium (https://taxonium.org/) view of the 1M-sample tree (A) before and (B) after optimization. The tips of the trees are colored based on the lineage assignments derived from a trained PangoLEARN model (30), with P.1 labels highlighted using red circles.

**Figure S3:**
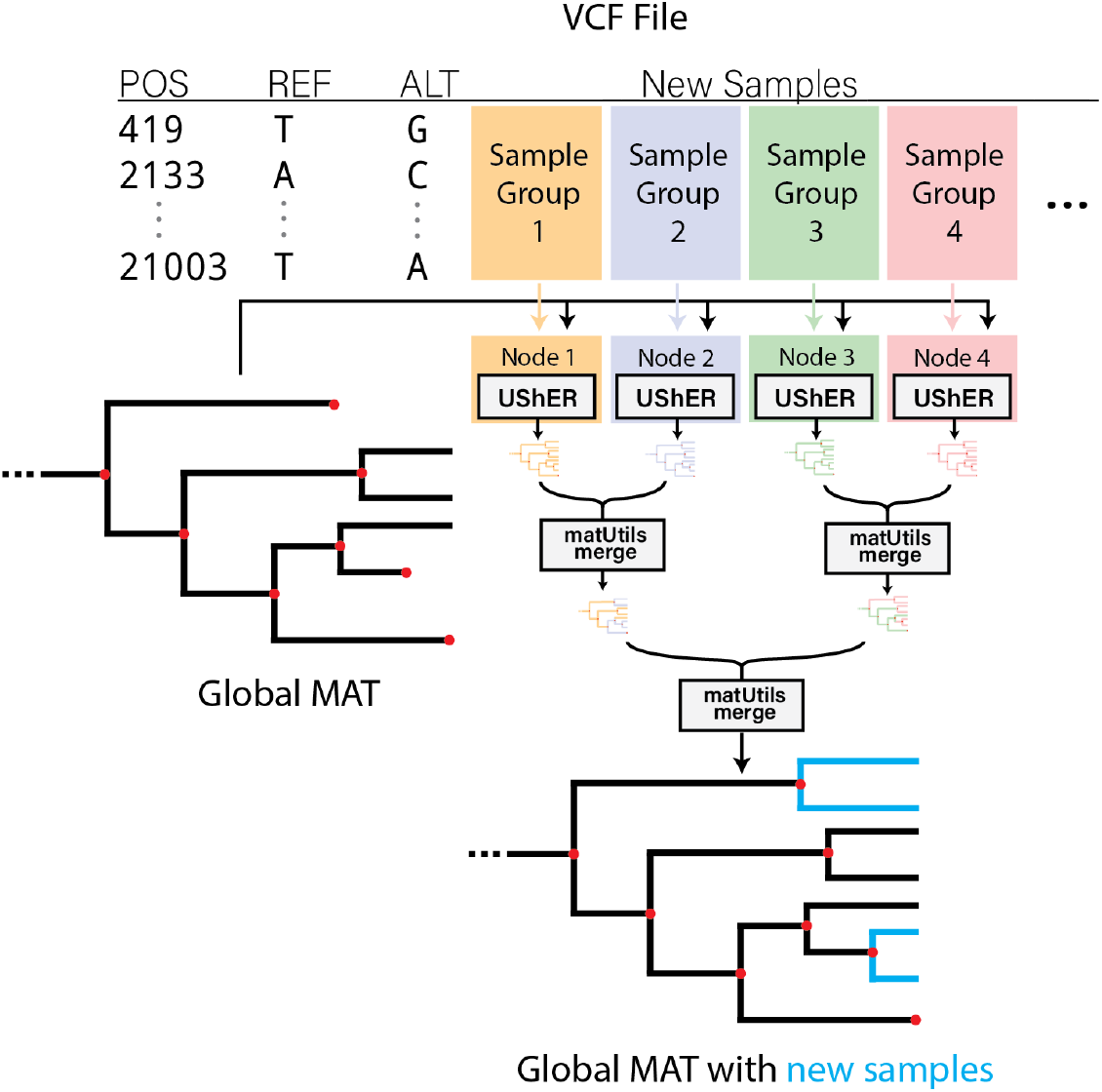
Multi-node parallelization of phylogenetic placement through UShER and matUtils merge.

**Figure S4:**
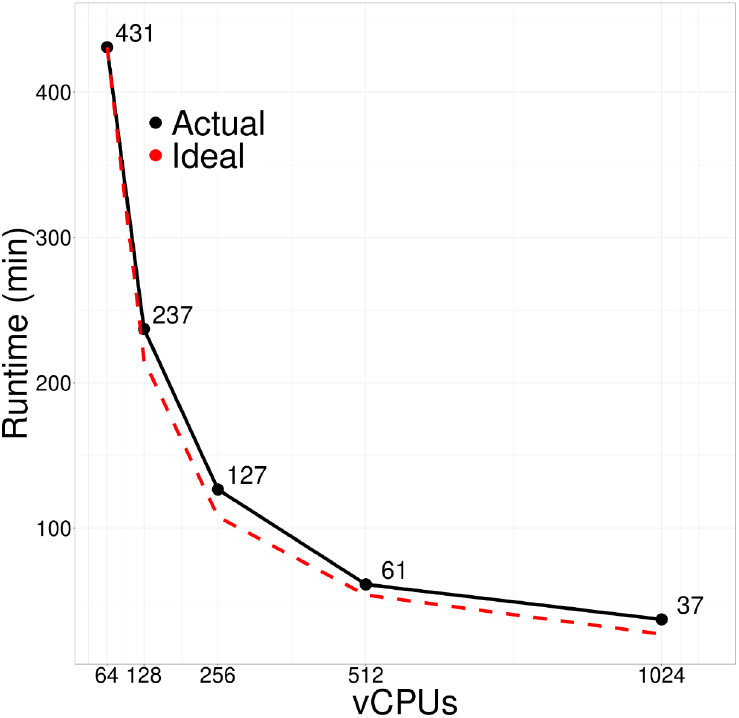
Strong multi-node scaling efficiency of UShER and matUtils merge. The numbers above each data point correspond to the actual runtime in minutes.

**Figure S5:**
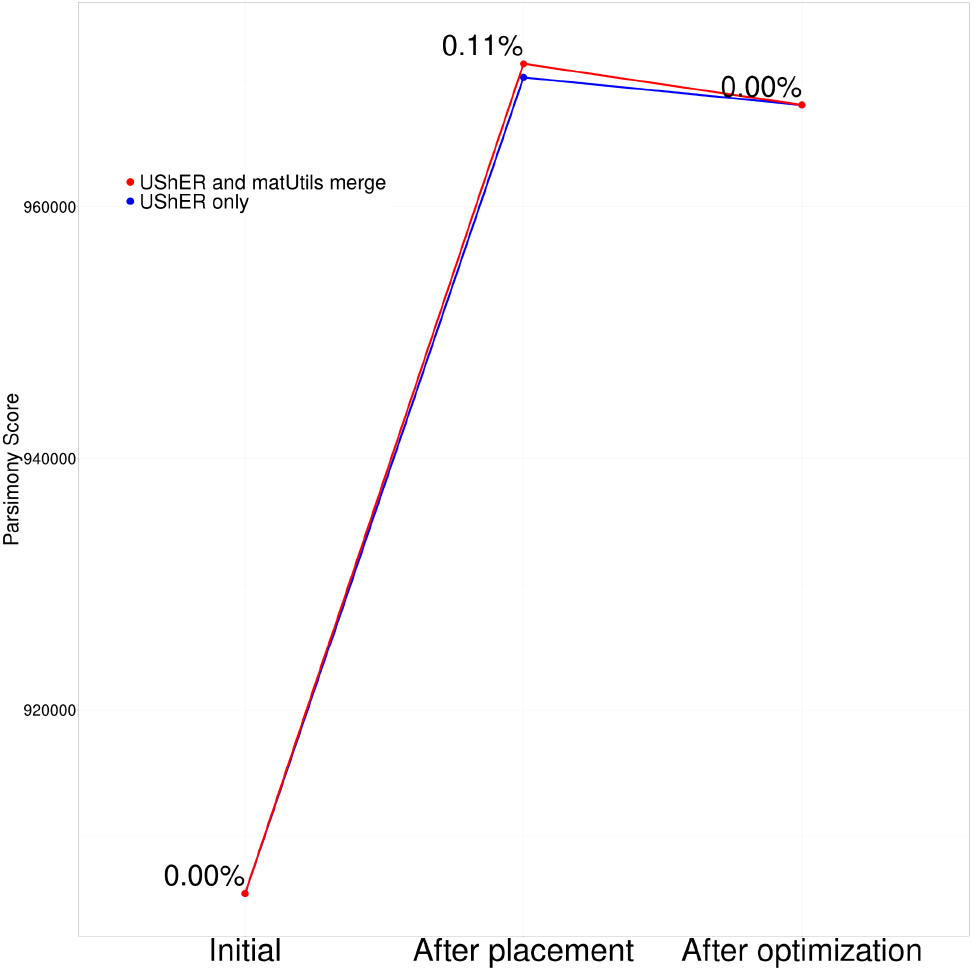
Parsimony scores starting from the 1M-sample tree (initial), after placement of 100K new samples in UShER and after optimization using matOptimize. The two lines correspond to the sequential incorporation of new samples in the placement step using UShER alone (blue) and the 16-way parallel incorporation using UShER followed by parallel merging using matUtils merge (red). The two approaches yield nearly the same final parsimony score after matOptimize, though the values are a bit different after the placement step. For matOptimize, we also provided as input, the VCF file containing the genotypes of all samples that allows it to maintain and re-resolve ambiguous bases at the phylogenetic tips.

**Table S1:** Parsimony score improvement with fixed radius and radius doubling optimization strategies in matOptimize.

| Tree size<br>(number of samples) | Parsimony Score Improvement |  |
| --- | --- | --- |
|  | Radius Doubling | Fixed Radius |
| 100K | 0.589% | 0.581% |
| 1M | 0.842% | 0.798% |
| 3M | 0.375% | 0.376% |

